# Monitoring free-living honeybee colonies in Germany: Insights into habitat preferences, survival rates, and Citizen Science reliability

**DOI:** 10.1101/2024.08.02.606354

**Authors:** Benjamin Rutschmann, Felix Remter, Sebastian Roth

**Affiliations:** BEEtree-Monitor, Munich, Germany; Department of Animal Ecology and Tropical Biology, Biocenter, University of Würzburg, Würzburg, Germany; Agroecology and Environment, Agroscope, Reckenholzstrasse 191, 8046, Zurich, Switzerland; Rachel Carson Center for Environmental Humanities, LMU Munich, Munich, Germany; Department of Science, Technology and Society (STS) at the TUM School of Social Sciences and Technology, TU Munich, Munich, Germany

**Keywords:** wildlife monitoring, *Apis mellifera*, urban ecology, wild pollinators, liminal species, wildlife conservation, wild-living honeybees

## Abstract

Our understanding of the western honeybee (*Apis mellifera*) predominantly stems from studies conducted within beekeeping environments, leaving the presence and characteristics of honeybees outside managed settings largely unexplored. This study examined the habitats, nesting sites, and survival rates of free-living colonies through personal monitoring of nest sites in Munich (N=107) and the coordination of Citizen Science monitoring across Germany (N=423). Within seven years we collected 2,555 observations on 530 colonies from 311 participants. Nesting preferences differed between urban, rural, and forested areas. Overall, we found that 31% of the occupied nest sites were in buildings and 63% in mature trees, with clear preferences for specific tree species. On average, only 12% of the personal monitored colonies in Munich survived annually, a figure that aligns well with other published studies but contrasts sharply with the significantly higher survival rates reported by Citizen Science (29%). We found that Citizen Science yielded significantly fewer updates per colony, underreported abandoned sites, and that 46% of overwintering reports overlapped with the swarming season and had to be excluded. To gain reliable survival data in Citizen Science projects, consistency and timing of reports need particular attention and regional swarming should be monitored too. This study enhances our understanding of the ecological dynamics, liminal state, and conservation needs of free-living honeybee cohorts, addresses potential monitoring biases, and suggests standardized data collection protocols for future monitoring projects.

## INTRODUCTION

The western honeybee (*Apis mellifera*) holds a significant place in the European entomofauna, facilitating the reproduction and genetic diversity of countless plant species, including many agricultural crops (Garibaldi et al., 2013; Breeze et al., 2014; Hung et al., 2018). As a cavity-dwelling species, *Apis mellifera* is adapted to live in forests, with tree hollows serving as its original nesting sites (Crane, 1999; Ruttner, 1988; Zander, 1949). Although the species has been used for honey harvesting since the Neolithic period (Crane, 1999) and plays a key role in modern commercial pollination services, it has undergone relatively little selective breeding compared to other similarly intensively managed animals (Oxley and Oldroyd, 2010). Nevertheless, among researchers and beekeepers alike, *Apis mellifera* tends to be perceived solely as a domesticated animal, found and researched under managed conditions. Consequently, most of our comprehensive understanding of the western honeybee as a species, its behaviour, and its ecology predominantly stems from research conducted with colonies under beekeeping conditions, while wild honeybee populations have been neglected in the modern apidological tradition (Stoeckhert, 1954; Kohl and Rutschmann, 2018; Seeley, 2019; Requier et al., 2019). The introduction of the parasite *Varroa destructor* (Anderson and Trueman, 2000) to Europe in the 1970s intensified this oversight, as the subsequent regular miticide treatment, suggested that only human-managed colonies could survive (Rosenkranz et al., 2010; Meixner et al., 2015).

Yet, this belies the fact that the western honeybee is also present outside the realms of beekeeping and human husbandry (Grindrod and Martin, 2021; Visick and Ratnieks, 2023a) and how little is known about the formation and dynamics of this cohort. Currently, stable wild honeybee populations within their original range are known to exist in Africa and the Southern Ural, and outside their original range in the Americas and Australia (Schneider and Blyther, 1988; Moritz et al., 2007; Ilyasov et al., 2015b; Ratnieks et al., 1991; Guzman-Novoa et al., 2024; Seeley, 2007; Bozek et al., 2018; Oldroyd et al., 1997; Chapman et al., 2008). Studies on these populations usually focus on habitats where managed and non-managed colonies are relatively separated, or where the density of wild colonies matches or surpasses that of managed colonies (Jaffé et al., 2010; Visick and Ratnieks, 2023a). The news about stable populations of western honeybee colonies outside beekeeping raised the interest of beekeepers in how to escape treatment (Seeley, 2019; Remter, 2021) and spurred considerable research in this field. Free-living colonies have been documented in various environments also in Europe, ranging from forests, to electric power poles in agricultural landscapes and rural and urban areas (Oleksa et al., 2013; Kohl and Rutschmann, 2018; Requier et al., 2020; Oberreiter et al., 2021; Kohl et al., 2022; Rutschmann et al., 2022; Hassett et al., 2018; Browne et al., 2020; Moro et al., 2021; Bila Dubaić et al., 2021; Lang et al., 2022; Visick and Ratnieks, 2023b; Cordillot, 2024). In Europe, however, the situation is different due to its fragmented landscape (Ibisch et al., 2016; Lesiv et al., 2019) and the high density of managed colonies (Phiri et al., 2022; Jones, 2004), which means there is no spatial and genetic barrier between managed and free-living colonies in most parts. The primary differences between free-living colonies and managed ones lie in their nest site ecology and their *modus vivendi*. Consequently, we propose referring to them as cohorts of local honeybee populations that are neither *fully wild* nor *domesticated* but exist in a *liminal state*.

To understand the free-living honeybee cohort more completely, it is crucial to examine not only their nesting sites but also the survival rates which are critical for assessing their genetic contribution to the local honeybee population. However, most of the studies on free-living honeybees in Europe have not systematically monitored individual colony survival, and have instead reported on nesting sites without comprehensive knowledge of the life histories of individual colonies– exceptions include Rutschmann et al. (2022), Kohl et al. (2022), Lang et al. (2022), Cordillot (2024). Due to their hidden locations in cavities high above the ground, free-living colonies lead secretive lives (Kohl and Rutschmann, 2018; Remter, 2021): finding them and repeatedly collecting data in numbers high enough for statistical inference is very time-consuming and requires specialized skills and equipment (Kohl and Rutschmann, 2018; Kohl et al., 2022). One approach that has garnered significant attention in the study of wild animals and biodiversity monitoring is the utilization of Citizen Science (Pocock et al., 2018; Fraisl et al., 2022; Koffler et al., 2021; Weissmann et al., 2023). Citizen Science offers the advantage of enlisting the help of many individuals who, in our case, shared the task of finding and monitoring colonies, thereby extending the geographic and temporal scope of the research beyond what researchers could achieve alone (Henneken et al., 2012; Lesiv et al., 2019; Hsing et al., 2022). Also, although previous studies have acknowledged the importance of Citizen Science in data collection (Moro et al., 2021; Bila Dubaić et al., 2021), none have yet investigated the quality of the data generated by Citizen Science and how to validate such reports.

With the BEEtree-Monitor, we developed a web-based monitoring scheme to study the habitats, nesting sites, and life histories of free-living honeybee colonies. Over a span of seven years (2016-2023), we collected various parameters of 530 nest sites together with longitudinal occupation data: 107 nest sites monitored by ourselves in the Munich region and 423 by citizen scientists mostly in Germany. Besides the main analysis we compare these two approaches to evaluate the validity and potential biases of Citizen Science reports and methodology. Through this study, we aim to provide a broader understanding of honeybees as a species that also exists outside human husbandry and offer guidelines to leverage future Citizen Science projects for effective monitoring of free-living honeybee colonies.

## METHODS

### Data collection and data curation

The monitoring and data collection process for this study was implemented through a combination of personal monitoring (PM) and reports by volunteering supporters with highly diverging skill levels and knowledge sets (citizen science monitoring, CS). Leveraging our own experiences in surveying free-living honeybee colonies and third-party reports, we developed an advanced monitoring scheme tailored to our target groups. Initially, participants were asked to fill in a designated form and send it to us via email (supplementary material ̶ SM). In 2018, we constructed an online platform specifically tailored for Citizen Science monitoring, launched as a website (BEEtree-Monitor; www.beetrees.org; see SM for further information). The CS recruiting was facilitated through social media outreach, presence in public media and beekeeping journals. The community was maintained via regular newsletters with guiding and motivating information. To enable as many volunteers as possible without offering a special training, our protocol focused on location, easily measurable nest site parameters, and continuous, repeated observations with specific date, time and focus (Figure 1A). Precise GPS coordinates allowed us to investigate the nesting and foraging habitats of the colonies, and the nest site parameters were used to analyse the swarms’ preferences for different nest types such as hollows in building structures (e.g., chimneys, window blind boxes or facade compound insulations) or hollows in different tree species. Additionally, we sought information on entrance directions and height, as well as the trunk diameter for trees. The online platform also featured an open-text/commentary field for participants to provide other relevant details observed.

**Figure 1:**
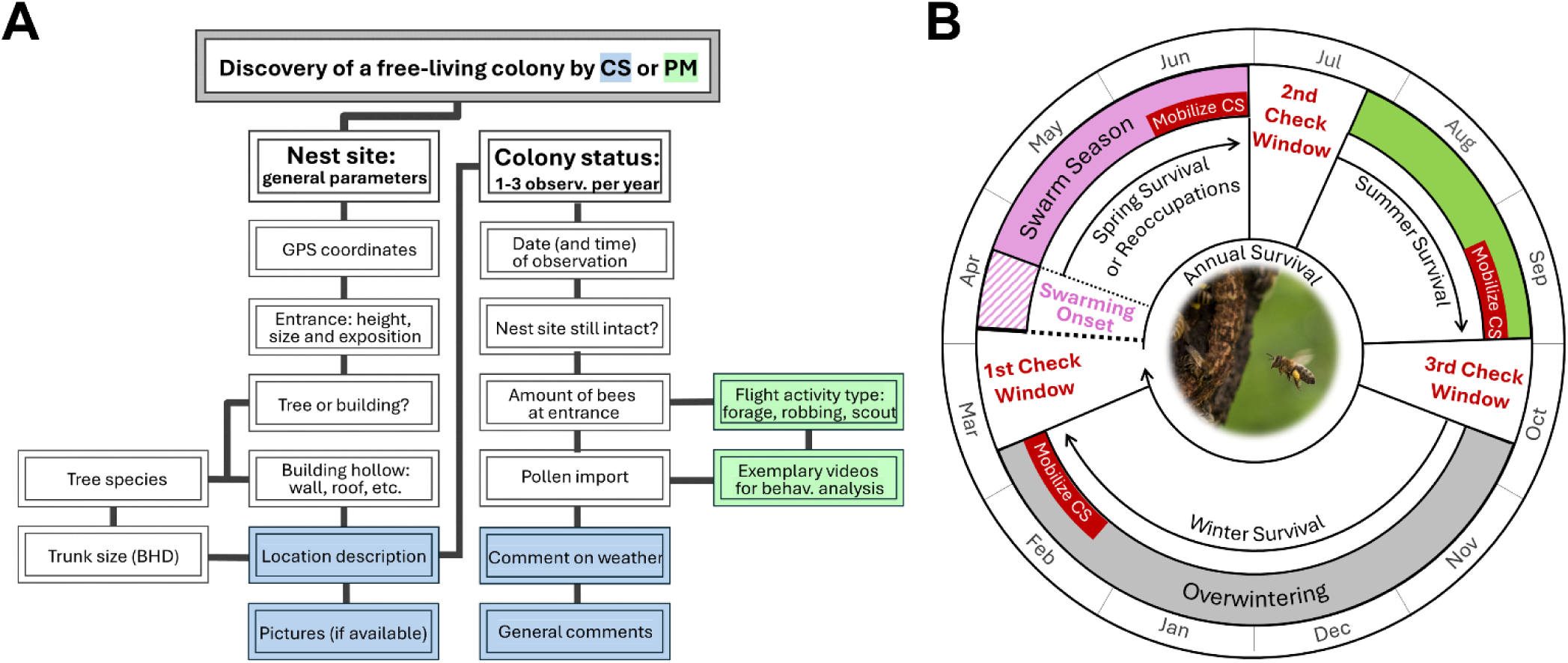
A) Monitoring protocol for Citizen Science (CS) and personal monitoring (PM) with steps for recording nest site parameters and colony status. Location descriptions, weather comments, general comments, and supplementary materials like pictures and videos helps validating the data in CS (indicated in light blue). In PM, we also collected information regarding distinct types of flight activity (indicated in light green). B) The annual monitoring schedule for free-living honeybee colonies outlines three critical observation windows: 1st check window (before the onset of the swarming season in late March and April), 2nd check window (post-swarming season in July), and 3rd check window (in autumn). These checks focus on different survival metrics: Winter survival, spring survival and reoccupations and summer survival, respectively. Citizen scientists (CS) mobilization should be done before each check window to ensure timely data collection. The yearly onset of swarming shifts and must be monitored closely to ensure accurate data on colony survival. The inlet picture shows pollen import into the colony an indication for brood production (photo: Felix Remter).

While both Personal Monitoring and Citizen Science follow the same basic protocol, their distinct nature allows for a comparison of the influence of monitoring type (Figure 1A):

> PM: This method entailed meticulous checks for visual signs of pollen import (indicating brood production, marking the colony as active) and frequent short-term observations of nest sites exhibiting ambiguous flight patterns or no observed foraging for pollen. We filmed some of the typical or ambiguous situations to collectively analyse and relate them to continuous observations at the site. This monitoring protocol substantiated our survival results, the classification of flight patterns and the ideal standard we aimed to achieve with CS.
>
> CS: This approach involved providing participants with protocols and guidance to e.g. prioritize pollen import observation. Accessible explanations in the online form and regular emails encouraged detailed reporting during key observation periods.

In total only 36 nesting sites were found by the authors (and 494 by citizen scientists), however 107 locations were actively monitored by us (and 423 by citizen scientists). Our primary analysis focused on colonies located in Germany and several nearby Central European countries, including Switzerland (N=9), Austria (N=3), Czechia (N=2), and Luxembourg (N=3). To mitigate the influence of markedly different environmental conditions that could confound our study’s findings, we excluded observations of ten free-living colonies from geographically distant countries like France, the UK, Italy, Spain, Norway, and Ukraine.

### Analysis of nesting and foraging habitats

To classify the nesting and foraging habitats of the reported free-living colonies, we imported the coordinates of each colony’s location into QGIS version 3.16.2 (QGIS Development Team, 2020) and performed intersections (using point layers for nesting habitats and 2 km buffers for foraging habitats) with the land cover classes from the CORINE Land Cover map 2018 (European Environment Agency, 2018). We grouped different land cover types together and quantified the proportional contributions of five major land cover types: urban areas, cropland, grassland, deciduous forest, and coniferous forest (see SM for further details). For part of the analysis these land cover types were further grouped into the three categories: Urban (urban areas), rural (cropland and grassland), and forest (deciduous and coniferous forests). In the case of the foraging habitat, we chose a radius of 2 km as approximately 80% of honeybee foraging occurs within this distance (Rutschmann et al., 2023). Moreover, the landscape within the 2 km scale has been shown to measurably influence honeybee colony performance, affecting factors such as foraging rate, colony growth, and winter survival (Steffan-Dewenter and Kuhn, 2003; Sponsler and Johnson, 2015; Rutschmann et al., 2022, 2023).

### Observation scheme for colony survival statistics

We defined survival as the instance of a cavity being occupied from summer (during or after the swarming season) until the following spring, before the next swarming season commenced (Figure 1B). One valid report of an active colony in spring before the start of the swarming season (1^st^ check) is proof for an overwintering survival. Additionally, we implemented and encouraged participants to conduct one or two more annual checks: after the end of swarming (2^nd^ check) and in Autumn (3^rd^ check). Post-swarming checks at all known cavities (including the recently unoccupied ones) served finding the new founder colonies for the further monitoring while the Autumn check (3^rd^ check) detects summer deaths. In our study, colonies that died in summer/autumn were attributed as perished to the survival statistics of the following year. For assessing overwintering, observations typically take place in March or early April where it is crucial that weather conditions are suitable for honeybee foraging (e.g. no rain and temperatures above 12° C. [Kevan and Baker, 1983]) but before swarming and therefore reoccupation of cavities. Consequently, observations without suitable weather conditions for honeybee foraging were excluded. We only considered primary data for our analysis, oral reports or in retrospect reports (of colonies living for "several years" in the same cavity) were considered hearsay and excluded. Destroyed nesting sites were also not considered in the survival statistics for the year of destruction. Additionally, to annual survival rates we also calculated spring, summer, and winter survival rates to compare with other studies (see SM for further information).

A potential pitfall of reporting mere flight activity at the entrance is that certain behaviours such as robbing, or the presence of scout bees may be erroneously interpreted as signs of a living colony by less experienced observers. Therefore pollen import into a colony was used as an indicator of brood production, hence designating it as an alive colony. Under certain conditions, even in the absence of visible pollen import, we still considered the colony to be alive. These criteria included:

1. <u>Observation of foraging flight patterns:</u> Colonies were classified as active if regular and/or directional flight activity was reported convincingly in the commentary section. These patterns are differentiated from non-foraging activities specifically observed in PM:

- Scout bees typically exhibit distinct behaviours such as taking time to land, thoroughly exploring the entrance before entering the cavity, performing slow orientation flights during departure, and might defend the entrance against other scouts when near swarming.
- Robbing bees display violent interactions with defenders if attempting to rob a living colony. If emptying a perished colonies stock, robbers initially perform orientation flights akin to scouts but more hectic with a distinct ‘bouncing off’ landing pattern.
2. <u>Reliability of observer comments:</u> If the observer was suspected to credibly discern differences in flight patterns indicative of foraging rather than robbing or scouting due to a competent comment given, the colony was considered active.
3. <u>Frequency and timing of observations:</u> Multiple observations made at short intervals that consistently indicated foraging behaviour, as opposed to scouting or robbing, supported the classification of a colony as active.

### Exemplary estimation of colony density

Estimating the density of free-living honeybee colonies poses challenges due to the likelihood of substantial underreporting. To address this, we concentrated our density estimation efforts on the city of Munich (further details can be found in the SM).

### Onset of regional swarming

In 2019, two of the authors (SR and FR) established a website across Germany and a hotline in Munich for the discovery of honeybee swarms and their potential capture. This enabled us to amass substantial data (N=376) on the initiation of swarming activity in the years from 2019 to 2023 within the geographic extent of this study. These years also represent the period during which most of the nest sites presented in this study were found (N=378 out of 530 colonies) and monitored (N=307 out of 350 life history reports). For each year, the first reported swarm served as a conservative estimate of the commencement of the swarming season. The onset of swarming varied over the five-year period. Specifically, swarming began on 17 April in 2019, 6 April in 2020, 8 May in 2021, 28 April in 2022, and 23 April in 2023. Hence, the time difference between the earliest and the latest recorded yearly swarming onset was more than one month (32 days), suggesting it should be taken into account when planning CS initiatives and survival analyses of free-living honeybees. Especially in years where swarming occurs unexpectedly early (e.g. year 2020). A small subset of the free-living colony reports in this study stem from years before 2019 in which we lacked empirical data on the beginning of the swarming season. Hence, we conservatively selected mid-April as the presumed start of swarming for these years (Henneken et al., 2012).

### Comparing survival rates of free-living honeybee colonies

To analyse colony survival statistics across different years in Germany we compared results from CS and PM and included two published studies –Kohl et al. (2022) and Lang et al. (2022). Each dataset employed a distinct monitoring approach (see SM for further details). To ascertain the influence of various predictors including monitoring types and year on the odds of colony survival, we compared several mixed-effects logistic regression models. These models were constructed using the ‘glmmTMB’ package in R (Magnusson et al., 2017). The model selection process was guided by the Akaike Information Criterion (AIC) and the ‘emmeans’ package (Lenth and Lenth, 2018) was used for post-hoc comparisons. Model predictions and confidence intervals were generated using the ‘ggeffects’ package (Lüdecke, 2018). Residuals of the models were inspected with ‘DHARMa’ package (Hartig and Hartig, 2017).

### Statistical analysis of directional and height preferences in cavities

To evaluate directional preferences of honeybees when selecting cavities, we recorded the entrance orientation of nesting sites occupied by free-living colonies in the eight cardinal and intercardinal directions for both tree cavities and cavities in buildings and assessed whether the distribution of orientations was non-random by employing the Rayleigh test for circular statistics using Directional package in R (Tsagris et al., 2016). The height of cavity entrances in trees and building structures was investigated with a non-parametric Mann-Whitney U Test.

Further information on statistical analysis regarding number of reports per colony, reported colony status and timing of the reports can be found in the supplementary material.

All statistical analyses were performed using R software (version 4.3.1; R Core Team, 2016). For data wrangling and graphical representation of the results, we utilized ‘tidyverse’, ‘ggplot2’, ‘patchwork’, ‘see’ and ‘ggpattern’ (Wickham, 2017, 2016; Pedersen, 2020; Lüdecke et al., 2021; FC et al., 2024).

## RESULTS

### Nesting and foraging habitat

The 311 participants (including the authors) provided 2,555 observations on 530 free-living colonies (Figure 2A). We found 57 percent of the reported colonies to dwell in urban areas with high human density, 14 percent in deciduous forest, 14 percent in cropland, 9 percent in grassland and 5 percent in coniferous forest (Figure 2B). The distribution of nesting habitats was significantly different from the proportional distribution based on land cover types in Germany (χ² = 99.81, df = 4, p < 0.001). Thereby, the proportional distribution assumes honeybees have no habitat preferences and their discovery is equally probable in all habitats. Colonies were disproportionately more often found in urban areas, equally often in deciduous forests, and less often in grassland, cropland and coniferous forests relative to the habitat availability in Germany. Calculating the available foraging habitat 2 kilometres around each nest site–where colonies are mainly operating to find their food–we found similar patterns. 41 percent of the available foraging habitat of the reported colonies were urban areas, 27 percent cropland, 12 percent deciduous forest, 11 percent grassland, and 9 percent coniferous forest (Figure 2C).

**Figure 2:**
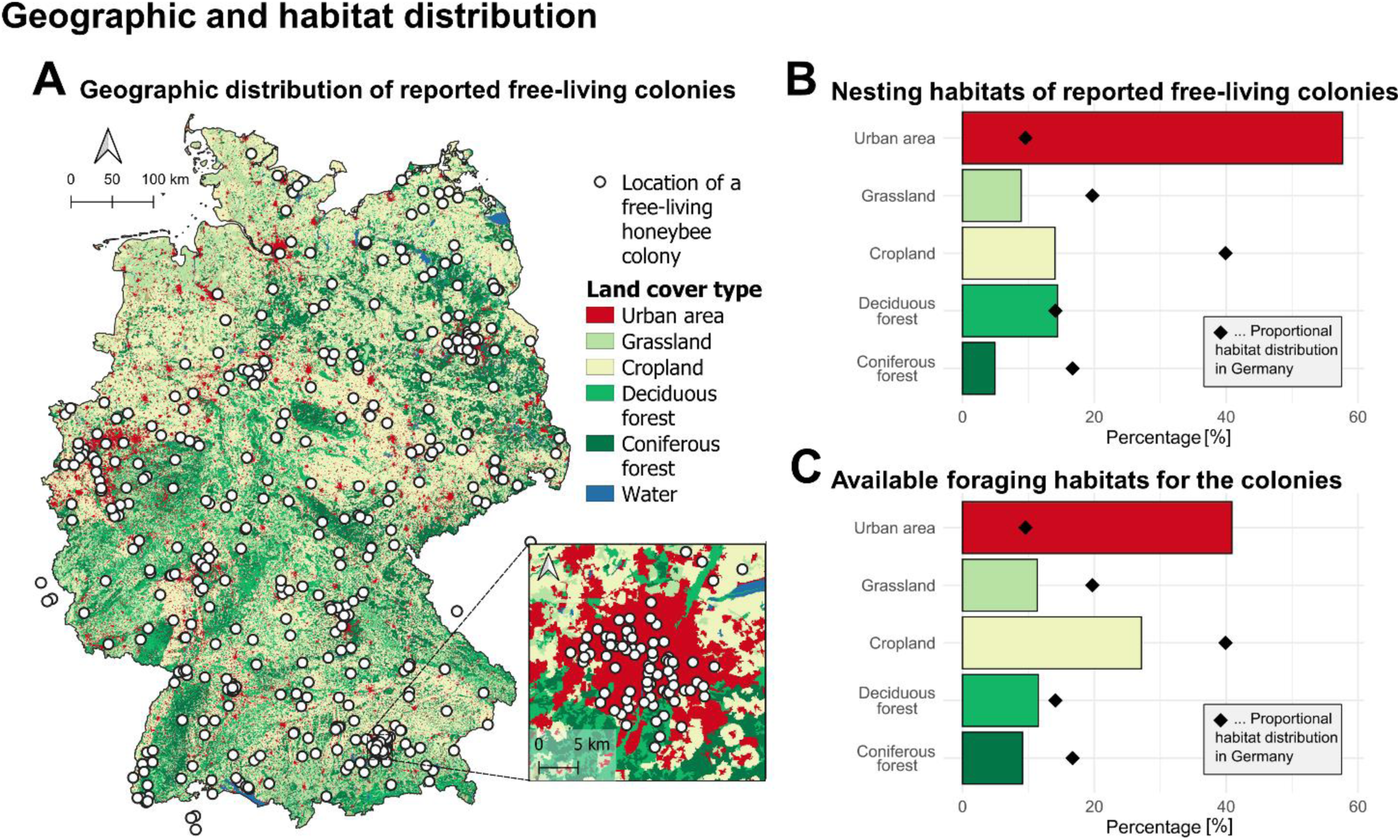
A) Geographic distribution of the free-living colonies reported, overlaid on the grouped main land cover types within Germany (CORINE Land Cover map 2018). The inlet shows the free-living colonies in the Munich region (3 colonies outside Germany are hidden by the inlet). B) Proportional distribution of nesting habitats where free-living colonies have been reported in comparison to the relative distribution of different land cover types across Germany depicted by black diamond symbols. C) Foraging habitats that is available to the bees (habitat within a 2 km radius around each colony). Again, black diamond symbols show the relative distribution of a land cover types across Germany.

### Nesting characteristics

The ratio of tree cavities to nesting sites in buildings, as well as the proportions of reported tree species, differed between urban, rural, and forest habitats. For the reported colonies in urban areas (57%, N=304), tree cavities comprised 52% of all cavities, while building cavities accounted for 40% (other cavity types: 8%; Figure 3A). In rural areas (23%, N=121), tree cavities were again dominating (68%), followed by cavities in buildings (21%). In forest areas (20%, N=105), tree cavities accounted for 81% of all nesting sites, while cavities in buildings represented 13%. Looking at the whole dataset 63% (N=324) of the colonies were found in trees, 31% (N=161) in building structures, and the remaining 6% (N=32) in other types such as rock crevices (N=3) and open nesting (N=15) (see figure in SM). Among tree species, Lime (*Tilia* spp.) was the most frequently occupied (N=59; 18%), followed by beech (*Fagus sylvatica*, N=45; 14%), oak (*Quercus* spp., N=41; 13%) and ash (*Fraxinus excelsior*, N=35; 11%) (Figure 3B). However, the distribution of these species differed across habitats (Figure 3B). Especially in urban areas, the diversity of tree species used by bees is much higher, with lime (23%), ash (*Fraxinus excelsior*, 16%), plane (*Platanus acerifolia*, 11%), and horse chestnut (*Aesculus hippocastanum*, 7%) being the most frequently occupied.

**Figure 3:**
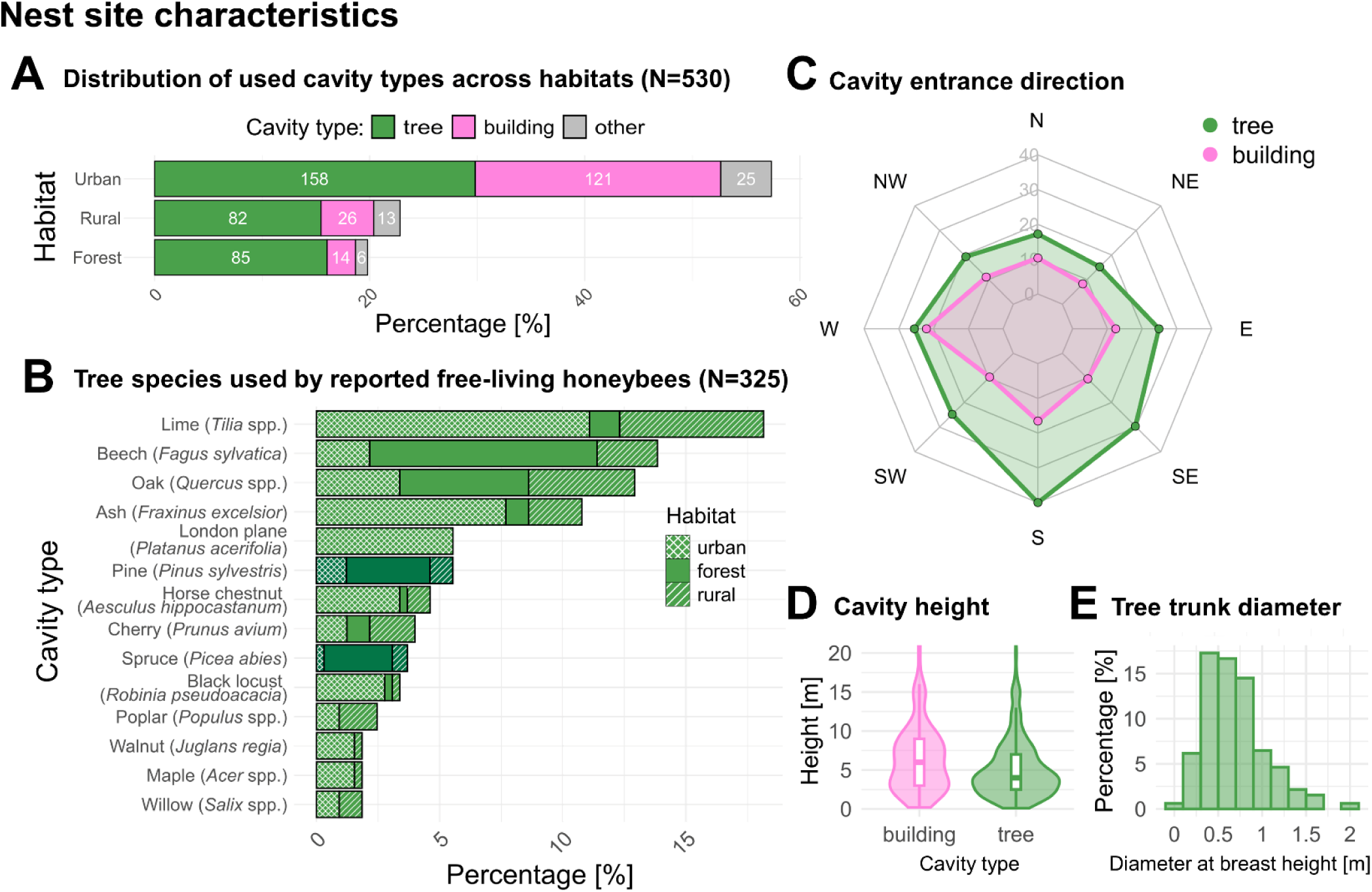
A) Proportional distribution of nesting cavity types (tree, building or other) across the habitats. While the x-axis gives the overall percentage, white numbers indicate the count (N) of trees, buildings, and other cavities. B) Distribution of tree species used as nesting sites in this study. Pattern fills illustrate the distribution across different habitats and darker green fill represents coniferous tree species. C) Preferred nesting entrance directions for cavities in trees and buildings. D) Cavity height distribution: A violin plot with an embedded boxplot that demonstrates the range and distribution of the heights of cavities occupied E) Distribution of tree trunk diameters (measured at breast height) for trees that hosted honeybee colonies.

Our study uncovered clear patterns in common flight entrance directions of nest sites used. For colonies situated in tree cavities, the observed frequency distribution across the eight cardinal and intercardinal directions was significantly non-uniform (Rayleigh Test Statistic = 20.23, Bootstrap p-value = 0.0010; N = 285). We found a significant preference for southern directions, with the highest frequency observed in the South direction (N = 58 or 20 %; Figure 3C). Similarly, for colonies located in buildings, the distribution was tested to be marginally non-uniform (Rayleigh Test Statistic = 4.85, Bootstrap p-value = 0.079; N = 146). Unlike tree cavities, cavities in human-built structures showed no clear directional preferences, although the West (N = 32 or 22%) and South (N = 24 or 16%) directions were observed most frequently (Figure 3C).

Our observations indicate that honeybee swarms predominantly choose nesting sites far from the ground (mean and median entrance height: 5.7 m and 4.5 m; Figure 3D). There was a statistically significant difference in the height of the cavity entrance in trees and in buildings (Mann-Whitney U Test W = 13482, p < 0.001). The entrance heights of cavities occupied in trees ranged from 0.1 to 30 meters, with a median of 4 meters. In contrast, the heights of cavities entrances in man-made structures ranged from 0.2 to 40 meters, with a median of 6 meters.

Additionally, we found that the diameter at breast height (DBH) of trees harbouring free-living honeybee colonies was on average 0.64 meters (median: 0.69 meters, Figure 3E), suggesting that colonies are dependent on trees with a substantial trunk diameter.

### Occupation rates and colony density across seasons in Munich

We investigated cavity occupation rates in the city of Munich during three distinct seasonal periods (spring, summer and autumn, see SM for further details). Multiplying the occupation rates with the density of 0.58 cavities per square kilometre that we know of in Munich (92 nest sites on an area of 160 km², see SM), we estimated the minimum density of free-living honeybee colonies in Munich to be approximately 0.06 colonies per square kilometre in spring, 0.42 in summer, and 0.28 in fall. It is important to note that the reported densities should be viewed as minimum estimates, given the likelihood that a significant number of colonies remain undetected and unreported. Based on these occupation rates, we infer that the number of colonies during summer is roughly seven-fold higher compared to spring. Conversely, the number of colonies in fall is approximately 33% lower than in summer.

### Survival rates and the impact of monitoring type and year

We analysed the life histories of 343 free-living honeybee colonies over the period from 2016 to 2023. Of these, 151 survival reports were provided by citizen scientists, while 192 were personally observed in Munich. It is important to note that a single colony could have had multiple survival reports, as it was monitored across different years. The yearly survival rates for the two monitoring types are depicted in Figure 4. In most years, survival rates reported by citizen scientists were higher than those observed through personal monitoring. Notably, in the spring of 2019, all PM colonies (N=32) perished, while 4 out of 17 colonies monitored through CS were reported as having survived. We observed a similar pattern when analyzing seasonal survival rates. For PM, survival rates were 87% in spring (N=23), 82% in summer (N=173), and dropped to 21% in winter (N=139). In contrast, CS survival rates were consistently higher, with 100% in spring (N=24), 93% in summer (N=180), and 46% in winter (N=98; see SM for more information).

**Figure 4:**
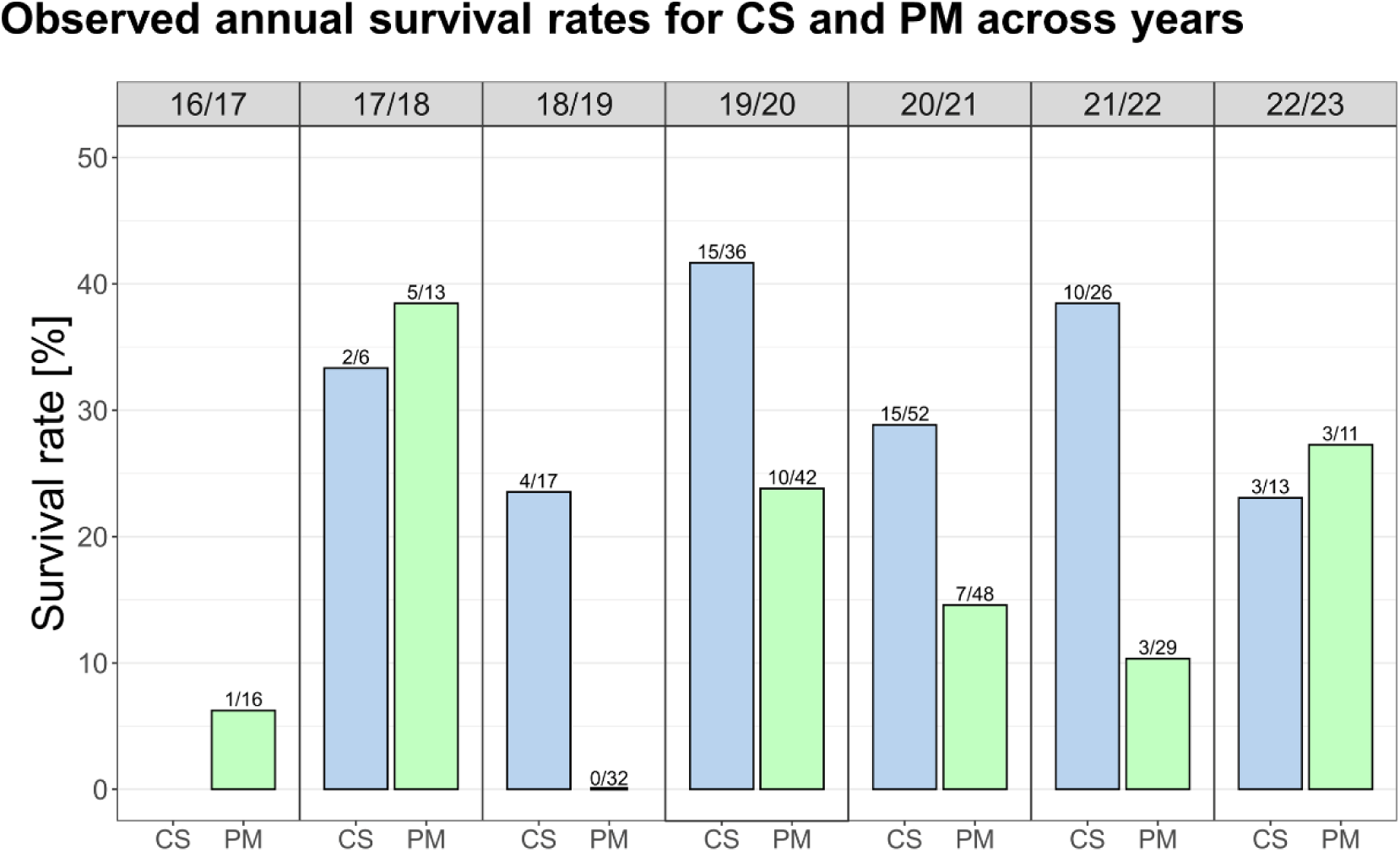
Comparison of observed colony survival rates from Citizen Science monitoring (CS, light blue) and personal monitoring in the Munich region (PM, light green) across the years. Numbers indicate the colonies that survived compared to the total reported.

Additionally, we consulted data from two published studies to determine whether the differences between CS and PM were likely due to biases in CS or actual differences. Kohl et al. (2022) looked at 112 colonies in German managed forest landscapes from 2017 to 2021, while Lang et al. (2022) investigated 30 colonies in Dortmund, Germany from 2018 to 2022 (Figure 5A)–more detailed information about the datasets and the reanalysis e.g. with the swarming timepoints observed in this study is provided in the SM. Model selection, guided by AIC values and convergence considerations, favoured including ‘year’ as a fixed effect alongside the ‘type of monitoring’ (CS, PM, Kohl et al. (2022) and Lang et al., (2022)). As a random effect ‘colony id’ was included to account for repeated measures on the same subjects across years. The likelihood ratio test (LRT) results from the glmmTMB model indicated that both ‘monitoring type’ (Chi-squared = 15.92, df = 3, p = 0.001) and ‘year’ (Chi-squared = 18.28, df = 6, p = 0.006) significantly contributed to the model. The estimated probability of survival was notably higher in CS (29%) than in PM (12%, p = 0.005) and in ‘Kohl et al. (2022)’ (13%, p = 0.02), but not significantly different from ‘Lang et al. (11%, 2022)’ (p = 0.32; probably due to the small number of colonies) (Figure 5B). While minor variations may arise from regional or habitat conditions, the significant discrepancies observed are likely due to reporting biases in CS compared to systematic surveys by experts.

**Figure 5:**
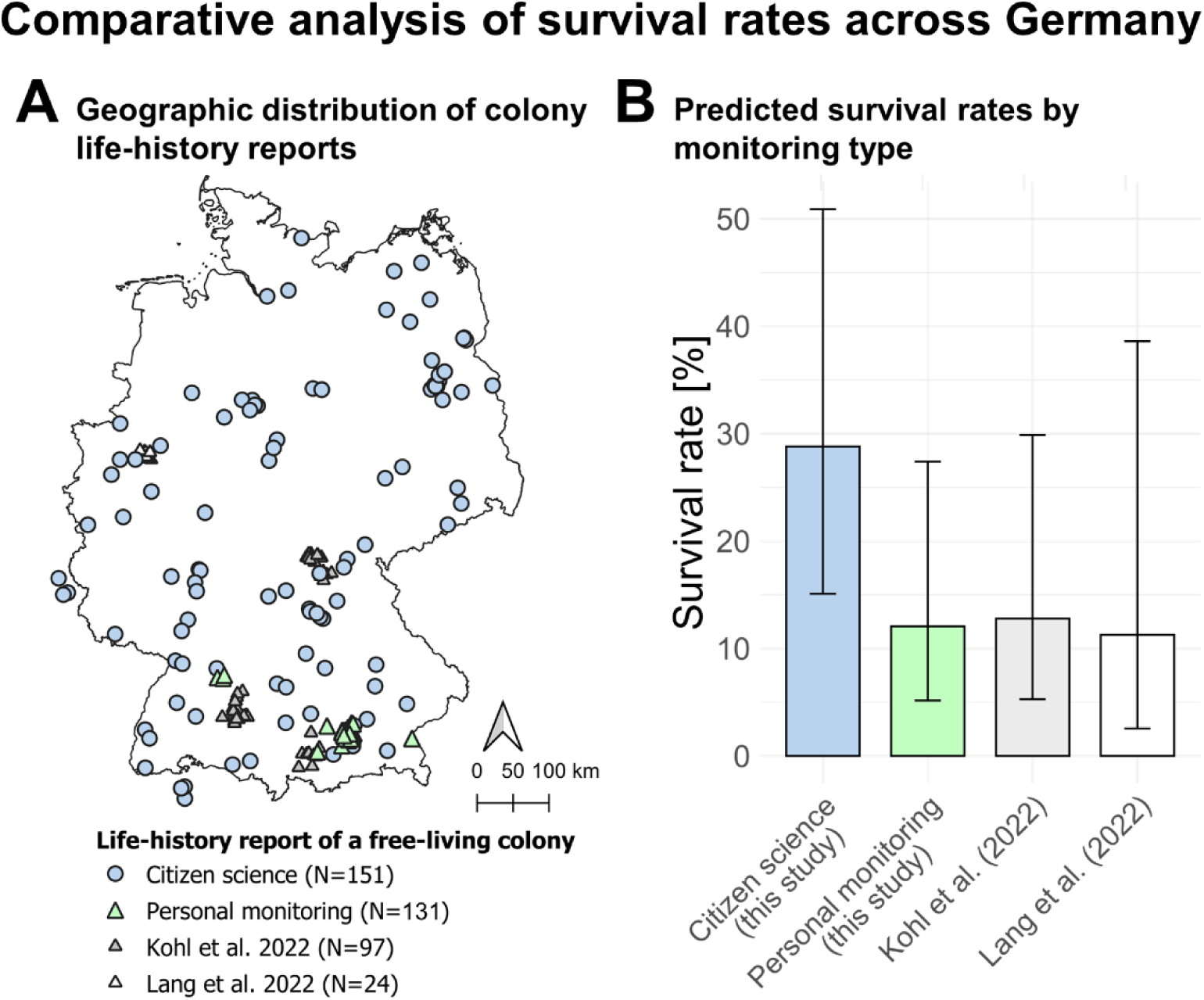
(A) Geographic distribution of life-history reports of free-living colonies from this study (CS and PM) and two published datasets –Kohl et al. (2022) and Lang et al. (2022). (B) Model estimates and 95% confidence intervals for average colony survival rates for the different monitoring types and studies.

### Biases with different monitoring types

Our dataset comprised 423 nest sites with 1,064 Citizen Science (CS) observations and 107 nest sites with 1,491 personal monitoring (PM) observations. We noted a significantly lower number of CS reports per colony compared to PM (Wilcoxon rank-sum test: W = 3950, p < 0.001; Figure 6A). Furthermore, we found a stark contrast in the distribution of "alive" versus "dead" colony status reports between CS and PM (Pearson’s Chi-squared test: χ2 = 176, df = 1, p < 0.001; Figure 6B), where 76% of CS reports indicated alive colonies compared to 42% in PM.

**Figure 6:**
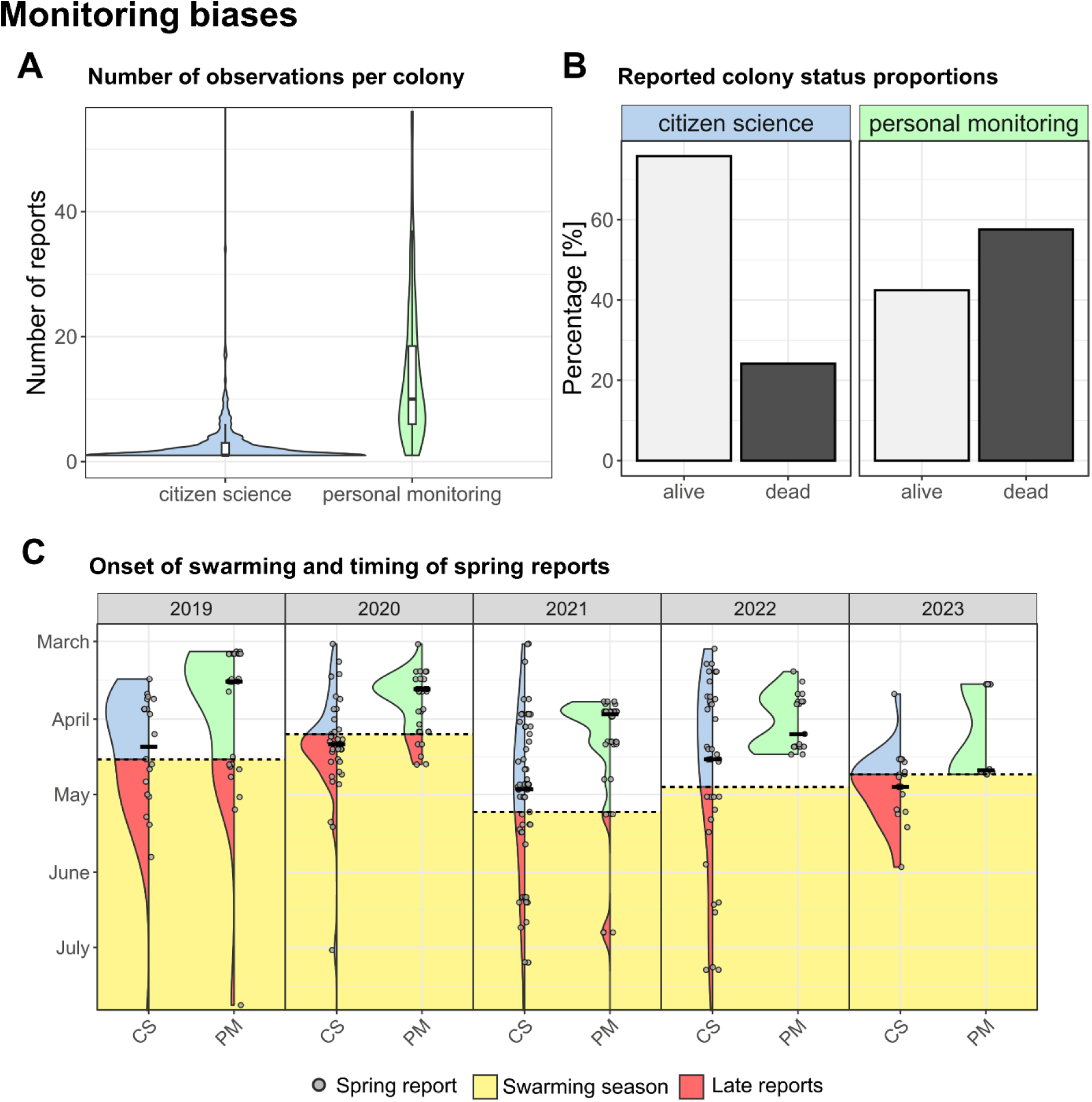
Analysis of reporting biases between Citizen Science monitoring (CS, light blue) and personal monitoring in the Munich region (PM, light green). A) Violin plot with included boxplot illustrating the distribution of the number of observations per colony for the whole monitoring timespan. B) Proportional representation of reported colony statuses (“alive” in light grey, “dead” in dark grey) for CS and PM. C) Temporal variation in colony statuses reported post-winter for the two different monitoring approaches in relation to the swarm season (in yellow) for the years 2019-2023. Note: The dashed line marks the first observed swarm for each year, after which spring reports on colony survival status may not be reliable anymore. Bold horizontal lines indicate median values of the timing of CS and PM reports.

To assess the effectiveness of the reporting in spring, we quantified the proportion of reports considered ‘late reports’, defined as submissions after the swarming start date for that year. The median reporting date after winter for CS was consistently later than for PM, specifically 21 April versus 29 March, respectively (mean: 3 April for PM vs. 2 May for CS) (Figure 6C). Consequently, a substantial proportion of CS reports were rendered unusable annually, i.e. around 46 % (84 out of 184) reports of CS could not be used each year, as they were reported too late (compared to 11% in PM, 15 out of 131 reports). These findings suggest an urgent need for improved reporting intensity and timelines within Citizen Science programs to ensure data reliability.

## DISCUSSION

To investigate the lives of free-living colonies and claims on high survival rates in Germany, we launched the BEEtree-Monitor project with a monitoring scheme designed to collectively gather data on free-living honeybee colonies. This initiative resulted in the most extensive database on the topic to our knowledge to date, enabling a comprehensive analysis of the nesting habits and life histories of these colonies over a seven-year period.

Deciduous forests are widely regarded as the original habitat of *Apis mellifera*, providing natural cavities in mature trees for nesting. Unlike most domesticated species, free-living honeybee colonies–when given the choice–continue to strongly prefer tree cavities, which are part of their original habitat. However, our data reveal that many of the tree species favored by honeybees are now rare in modern forests and are more commonly found in urban areas, along streets, in parks, or in cemeteries. The most common tree species by area in German forest environments are spruce (*Picea abies*, 25%), pine (*Pinus sylvestris*, 22%), beech (*Fagus sylvatica*, 15%), and oak (*Quercus* spp., 10%), collectively representing more than 70% of the trees (Schmitz, 2014). When examining tree species that house free-living honeybee colonies in German forests, these four species were also the most common, but the distribution was significantly different. While the deciduous species beech (35%) and oak (20%) were highly overused by bees, colonies were greatly underrepresented in pine (13%) and spruce (11%), the two most important coniferous tree species in managed forests. Although coniferous species are still predominant in managed forests, they do not provide as many suitable cavities for bees and other animals (Requier et al., 2020). Lime trees (*Tilia* spp.), which are of minor importance in managed forests, dominate both the urban and rural landscapes as alley trees and are heavily chosen by honeybees as nesting sites and sources of food (Tofilski and Oleksa, 2013). Our observations that ash trees are disproportionately favoured as nesting sites, raises concerns in light of the ongoing threat of *Hymenoscyphus fraxineus*, the causal agent of ash dieback across Europe. In Munich, nearly half of the inhabited trees are ash, and with over half of the city’s free-living colonies residing in buildings, the ash dieback could significantly increase human contact with these colonies. Urban planning should take into account the potential increase in human-wildlife interactions, such as those involving woodpeckers, pigeons, wasps, or hornets, and incorporate strategies to minimize conflicts. This could include offering designated cavities or nest boxes for different animals, as well as promoting habitats for beneficial species like solitary bees.

When examining the available foraging habitat around the reported colonies, our findings indicate that today’s forests play a minor role, area wise and because German forests do not provide diverse and rich foraging opportunities throughout the year (Rutschmann et al., 2023). Our calculations, however, indicate a notably high density of free-living honeybee colonies in urban areas, exemplified by Munich in summer (0.42 colonies/km²), surpassing those in managed German woodlands (0.23 colonies/km²), Polish rural areas (0.1 colonies/km²), and agricultural landscapes in Spain (0.2 colonies/km²) (Kohl and Rutschmann, 2018; Kohl et al., 2022; Oleksa et al., 2013; Rutschmann et al., 2022). Urban areas are less contaminated by pesticides and offer an array of potential nesting sites, such as cavities in buildings and mature trees in parks and gardens (Angold et al., 2006; Threlfall and Kendal, 2018; Gathof et al., 2022). Additionally, the floral diversity of urban areas provide an attractive foraging environment for both managed and free-living honeybee cohorts (Ayers and Rehan, 2021; Baldock, 2020; Young et al., 2021; Garbuzov et al., 2015; Samuelson et al., 2021).

This accumulation of free-living colonies in Munich and other cities could be attributed to several factors, including those mentioned above. It is also known that higher human population density correlates with higher densities of beekeepers, hives and consequently more escaping swarms searching for cavities (Oré Barrios et al., 2017; von Büren et al., 2019). For example, in Munich, the known density of managed hives is over 12 per km², surpassing the German average density by a factor of 4.3 (personal information from the veterinary office and German Ministry of Food and Agriculture, 2022). However, the proximity of honeybee colonies to humans in urban areas increases the likelihood of discovery, potentially introducing biases in monitoring. The dominance of reports from urban areas may skew perceptions of the distribution and density of free-living colonies across different habitats. This issue can be addressed by incorporating systematic approaches with random sampling techniques in future studies, such as beelining (Seeley, 2016; Kohl and Rutschmann, 2018; Radcliffe and Seeley, 2018; Chakuya et al., 2022).

Our findings reinforce the understanding that honeybees generally prefer elevated cavities for nesting (Seeley and Morse, 1978; Seeley, 2019). The observed significant difference in the height of nests between building and tree cavities may partially reflect a bias, where colonies in buildings are more likely to be observed despite higher elevation from ground level. Also, our study underscores the importance of mature, old-growth trees in providing suitable nesting sites for free-living honeybees (Bütler et al., 2013; Requier et al., 2020; Visick and Ratnieks, 2023b; Lindenmayer et al., 2012; Saunders et al., 2021). With an average diameter at breast height (DBH) of 64 centimetres, these large trees in our study proof to be vital for supporting diverse ecological communities. An interesting aspect is that free-living honeybees exhibited a pronounced directional preference for southern or southwestern exposures when selecting cavities. This preference aligns with the thermoregulatory benefits of southern orientations, which facilitate sun exposure and warmth, particularly beneficial during spring when colonies are emerging from winter. This finding is consistent with some previous studies that noted similar directional preferences (Seeley and Morse, 1978; Avitabile et al., 1978; Radcliffe and Seeley, 2018; Cordillot, 2024), though it contrasts with others that reported a random distribution of entrance directions (Seeley and Morse, 1976; Gambino et al., 1990). Additionally, in both cities and forests, a high proportion of bee-used hollows in trees and facade insulations are created by woodpeckers. While we cannot exclude an underlying preference of woodpeckers for certain directions and heights, the consistent directional preference observed in our study suggests a genuine selection by the bees themselves.

High managed honeybee densities in many European regions have sparked ongoing debates about food competition and pathogen transfer among pollinators (Geldmann and González-Varo, 2018; Alaux et al., 2019; Ropars et al., 2019; Saunders et al., 2018; Herrera, 2020; Iwasaki and Hogendoorn, 2021; Ghazoul, 2005; Casanelles-Abella and Moretti, 2022; Weissmann et al., 2023; Egerer and Kowarik, 2020). Such concerns could be particularly pronounced in urban settings where endangered solitary bees are present and the density of managed honeybee colonies is already high (Mallinger et al., 2017; but see Harder and Miksha, 2022; Steffan-Dewenter and Tscharntke, 2000). Yet, our findings suggest that the density of free-living colonies in Munich remains relatively low, at approximately 4% of the density of registered colonies in the city (see SM for more information). While the local densities of managed colonies require careful consideration, free-living colonies should not be considered problematic for urban pollinators nor for managed colonies from urban beekeepers ̶ see Kohl et al., (2023a) for an investigation on the parasite loads of free-living colonies.

To assess the self-sustainability of a population is important for the conservation of species with a clear status of being “wild” or “feral”. For some species as for the western honeybee the situation in Europe is more complex. Local European honeybee populations consist of deeply entangled cohorts of managed and free-living colonies in varying proportions. For centuries, the breeding of the former and the natural selection of the latter have impacted the overall population. With the advent of *Varroa destructor* and the continuing use of acaricides in most beekeeping practices, the ratio might have shifted in some European regions, including Germany. Yet, the species remains in a liminal state, and depending on future conservation strategies, populations might either shift towards full dependency on human care or retain and even enhance their capability to self-sustain by naturally adapting to new environmental circumstances.

In regions such as Spain and the UK survival rates of free-living honeybee cohorts suggest potential self-sustainability or near self-sustaining populations (Rutschmann et al., 2022; Visick and Ratnieks, 2023b). Notably, in Gwynedd, Wales, the use of acaricides in beekeeping is no longer necessary (Remter, 2015). Our personal monitoring analysis in Munich, however, indicates that the survival threshold required for self-sustaining cohorts — approximately one-third annual survival (Kohl et al., 2022; Rutschmann et al., 2024) — is not being met in Germany. This finding aligns with other studies conducted in variable habitats in Germany and neighbouring countries (Kohl et al., 2022; Lang et al., 2022; Cordillot, 2024), yet contrasts sharply with anecdotal claims from lay observers who report continuous and prolonged occupancy of cavities. Yet, these observations do not necessarily confirm extended lifespans of individual colonies; rather, it seems more plausible that many free-living colonies in Germany are recent escapees from managed apiaries, and that monitoring has not been conducted rigorously enough.

The factors contributing to the decline of free-living colonies are varied, encompassing both ecological and evolutionary aspects. From an ecological perspective, challenges such as a shortage of floral resources and lack of suitable and safe nesting sites can significantly impact colony viability (Kohl et al., 2023b; Rutschmann et al., 2023), while parasites, though present, may not play a major role in colony mortality in Germany at the moment (Kohl et al., 2023a). Another aspect to consider is the evolutionary impact of modern beekeeping practices: They focus on breeding for desirable traits such as maximized honey production, reduced swarming tendencies, and docility (Seeley, 2019). Breeding efforts for varroa resistance exist, but they have not yet made a significant impact (Guichard et al., 2023), and the continuous medical treatment of managed colonies prevents tolerance traits from prevailing at the population level (Blacquière et al., 2019; Neumann and Blacquière, 2017). Additionally in Germany, the replacement of the native subspecies *Apis mellifera mellifera* with non-native subspecies may have profound implications on the genetic diversity and adaptability of honeybee populations in this area, potentially influencing their ability to thrive independently (Büchler et al., 2014; Dustmann and others, 1988; Meixner et al., 2015). The ongoing decline in genetic diversity and introgression of non-native subspecies further threaten the resilience of free-living colonies (Espregueira Themudo et al., 2020; Muñoz and De la Rúa, 2021). Understanding the complex interplay of these factors is crucial for developing reliable conservation strategies for free-living cohorts.

The issue of defining species as “wild” and self-sustaining is significant, especially as organizations like the IUCN assess species status based on these criteria. Emphasizing wilderness and self-sustainability can hinder conservation efforts for liminal species in danger of losing their ability to self-sustain. We suggest understanding honeybees in Europe as a liminal species, with larger cohorts living under beekeeping conditions and smaller ones living independently, both within a broad and shared geographic range. These cohorts are unlikely to be found independently of each other and are deeply interconnected through genetic exchange via mating and annual swarming. The survival rates of free-living cohorts influence the overall genetic diversity of honeybee populations in Europe, including the managed cohorts. However, even the environments shared by these cohorts do not align neatly with the modern nature/culture dichotomy. As mentioned earlier, most German forests are not “natural” wilderness spaces, while urban landscapes have, from a conservation perspective, become essential refuges for species considered “wild.” This suggests that no pristine habitat, in the sense of a "natural" environment, serves as a precondition for self-sustaining “wild” honeybee populations. Consequently, the traditional dichotomies of natural/artificial or wild/domestic may be misleading as categorical foundations for contemporary research and conservation efforts.

While the higher number of reports for colonies monitored by personal monitoring (PM) is partly due to the density of observations applied in the PM to better understand the flight patterns of robbing, scouting or survivor bees, there remains a concern with Citizen Science (CS) as many colonies in the CS dataset were reported only once and could not be used for the survival analysis. In fact, it is plausible that citizen scientists primarily report on active, living colonies, and may discontinue monitoring once a colony is no longer present, thereby creating a biased dataset. In our study, approximately 76% of reports from citizen scientists indicated the presence of living colonies, in contrast to only about 42% in the PM dataset. This difference aligns with the 52% and 43% living colonies reported in two expert-monitored studies by Rutschmann et al. (2022) and Kohl et al. (2022), respectively. This discrepancy between reports on occupied and unoccupied cavities suggests that the ratio of reported colony status indicate inherent biases within the monitoring methods of layperson in contrast to experts. Understanding and addressing these biases is essential for accurately interpreting survival rates of free-living honeybees based on Citizen Science data. Our analysis also revealed a significant delay in reporting by citizen scientists in spring. This delay rendered a significant proportion of the data unusable for survival analysis. Despite accounting for this factor, the survival rates reported in CS data remained much higher, suggesting a likely bias. Our model predictions, incorporating monitoring type and data from CS, PM, and two published datasets, also point in this direction, reinforcing the potential bias introduced by CS reporting practices. Overall, the findings suggest that survival results from CS should be interpreted with caution.

A key challenge of our study, similar to many Citizen Science projects, is its serendipitous nature. While Citizen Science is extremely valuable for identifying the locations of free-living honeybee colonies across huge geographic scales, it presents complexities and significant efforts when it comes to monitoring these colonies coherently over time. Based on our experience in the first years of the study we optimized the PM monitoring for a better applicability in CS. To maximize the contribution of CS to bee research, we recommend adopting a standardized monitoring scheme (Figure 1B). This approach can be built on the successful protocol employed in our personal monitoring, which consisted of at least three site visits per year to detect pollen import (or typologized flight-patterns) and empirical data on the swarming season (especially its onset). The first visit should occur before the swarming season begins, the second visit after the swarming season to confirm newly established colonies and to include known but previously unoccupied sites to find new founder colonies, and the third visit in autumn to assess summer-deaths and the number of colonies going into winter. Citizen scientists need to be reminded early and often enough and briefed on appropriate timing and weather conditions for observations. Given the average height of the entrances (5.7 m), they should be equipped with sufficiently powerful binoculars or spotting scopes to reliably observe pollen import at each of the three visits. Additionally, observers could be trained to differentiate between various flight patterns, such as foraging, robbing, or scouting along the criteria offered in this study. Including a commentary section in the monitoring forms proved valuable, as it allowed observers to describe their observations in detail. This additional information helped us assess the reliability of the data, especially in cases where the data seemed suspicious or lacked information on pollen import. While we could then decide whether to include or exclude such reports in the final dataset, having consistent reports on pollen import is much more time-efficient and should be prioritized. In essence, structured and standardized monitoring projects are indispensable for thoroughly understanding the mechanisms underlying survival of free-living colonies. Long-term monitoring and more extensive geographic coverage would enhance our understanding of the survival and reproductive success of free-living honeybee colonies in Europe. Collaborative efforts combining scientific research with Citizen Science initiatives could prove beneficial in achieving these objectives. In this sense, overwintering survival serves as a practical and easily recordable indicator for identifying regions in Europe with high survival rates and enlarged free-living cohorts. Such an approach requires minimal investment in monitoring and does not rely on sophisticated kinship analysis, making it accessible for widespread Citizen Science participation.

## CONCLUSION

Given the growing trend of citizen science-based knowledge production, particularly for monitoring free-living honeybee colonies, we critically compared the citizen science dataset with data collected through our personal monitoring efforts. We identified epistemic pitfalls in the collection and interpretation of citizen science data and outlined potential strategies to optimize its use for further monitoring purposes. Despite a reporting bias towards more densely populated areas, the combined datasets provide a reliable and comprehensive foundation for assessing nest site parameters of unmanaged colonies across various land cover types in Germany. Importantly, the “low-tech” approach of our personal monitoring offered a valid foundation for survival analysis. Our findings contribute to filling a gap in our knowledge about the western honeybee and enhance our understanding of its free-living cohort in Germany. Given the low survival rates of free-living colonies in Germany compared to other regions in Europe and abroad, more detailed and comparative research is needed to develop conservation strategies for free-living cohorts. It is clear already that, conservation efforts should prioritize the preservation of old-growth forests and the protection of long-lived tree species across all habitats to maintain crucial nesting habitats for many species including free-living honeybees.

## ACKNOWLEDGMENTS

We express our gratitude to the BEEtree-Monitor community and all the citizen scientists who provided data for this study. A list of data contributors and those who helped shape the monitoring protocol can be found in the Acknowledgment Appendix in the SM.

## AUTHORS’ CONTRIBUTIONS

Conceptualization: SR-BR-FR

Methodology: SR-FR-BR

Software: SR

Formal analysis: BR

Investigation: SR-FR Resources: FR-SR-BR

Data curation: SR-BR-FR

Writing - original draft: BR-FR

Writing - Review & editing: SR

Visualization: BR

Administration: SR-FR

## DATA AVAILABILITY STATEMENT

Data used in the submitted manuscript can be made available to a limited degree after a reasonable request to one of the corresponding authors. Personal data of citizen scientists will not be available due to violation of privacy.

## ETHICS APPROVAL

No honeybees were harmed during this study, which used non-invasive monitoring methods. Ethical approval was not required as honeybees are invertebrates.

