## Supplementary Information for "Monitoring free-living honeybee colonies in Germany: Insights into habitat preferences, survival rates, and Citizen Science reliability"

### ACKNOWLEDGMENT APPENDIX

We thank Andrea Voit, Andreas Schmitt, Andrej Barth, Anja Salg, Anna Dünser, Antonio Gurliaccio, C. Hutschenreuther, Christian Ahrens, Claudia Schneider-Ludwig, Frank Schuberdt, Frank Soukup, Gottfried Schumann, Hagen Morscheck, Hannes Oberreiter, Ingo Schüler, Jens Eickmeier V.N., Kai Oliver Siegenthaler, M. Billmeier, Markus Kimpel, Martin Dannenberg, Michael Ostheer, Moses Martin Mrohs, Olivia Ortlieb, Omar Alejandro Machado Taboada, Peter Pfanzelt, Sergej Hübert, Sven Büchner, Thomas Mull, Thomas Schneider-Ludwig, Vivien Otto, W. Schwarz, Waldemar Schmidt and Wolfgang Mück and many more who didn’t answer to our call for mentions in the acknowledgement. We would like to express our gratitude to the HOBOS Team of the University of Würzburg for their support with the BEEtrees project during the initial stages of our research. Special thanks go to Jürgen Tautz, Marina Kretzschmar, Gerhard Vonend, Kristina Vonend, Anneli Kiessling, Konrad Öchsner, Hartmut Vierle and the rest of the team. We also thank André Wermelinger, Uwe Lang, Frank Krumm and Patrick Kohl for their valuable input during the initial discussions on the monitoring protocol. Additionally, we acknowledge Valerie Kantelberg, Sigrun Mittl, and Andreas Schierling for their insightful discussions on the topic.
